# NGS-PrimerPlex: high-throughput primer design for multiplex polymerase chain reactions

**DOI:** 10.1101/2020.06.29.176834

**Authors:** Andrey Kechin, Viktoria Borobova, Ulyana Boyarskikh, Evgeniy Khrapov, Sergey Subbotin, Maxim Filipenko

## Abstract

**Summary:** Multiplex PCR has multiple applications in molecular biology, including developing new targeted NGS-panels. We present NGS-PrimerPlex, an efficient and versatile command-line application that designs primers for different refined types of amplicon-based genome target enrichment. It supports nested and anchored multiplex PCR, redistribution among multiplex reactions of primers constructed earlier and extension of existing NGS-panels. The primer design process takes into consideration formation of secondary structures, non-target amplicons between all primers of a pool, primers and high-frequent genome SNPs overlapping. Moreover, users of NGS-PrimerPlex are free from manually defining input genome regions, because it can be done automatically from a list of genes or their parts like exon or codon numbers. Using the program, NGS-panel for sequencing the *LRRK2* gene coding regions was created, and 354 DNA samples were studied successfully with median coverage of 97.4% of target regions by at least 30 reads.

**Availability and implementation:** NGS-PrimerPlex is open-source and freely available at https://github.com/aakechin/NGS-PrimerPlex/.

**Supplementary information:** Supplementary data

## 1 Introduction

Multiplex PCR is a fundamental approach to get many several fragment copies of single DNA molecule simultaneously. It has a lot of applications in different science areas, from kinship determination (Grover et al., 2017) to NGS library preparation (Cohen et al., 2018; Ermolenko et al., 2015). A transition from monoplex to multiplex reactions requires consideration of many factors that may influence PCR efficiency. It includes secondary structure formation by oligonucleotides and non-target hybridization, elongation and amplification with primers from different pairs. All of this should be taken into account during manual primer design.

To facilitate primer design process, many tools have been developed (**Supplementary Table 1**). However, none of them can completely simplify the process of developing new amplicon-based NGS panels. This library preparation approach has become widespread among both researchers and commercial companies due to its simplicity of utilization by end user and highly efficient target enrichment for DNA samples of different quality. This approach has become particularly relevant for the molecular genetic testing of tumors, where many target genes should be studied. At the same time, targets differ not only from one type of cancer to another, but also between patients with one type of oncology.

Here we describe a new tool for automatic primer design, particularly for creating new amplicon-based targeted NGS-panels, NGS-PrimerPlex. This program has many features that are useful during developing new NGS-panels (**Supplementary Table 1**). Among them we note the following features: (1) choosing genome regions based on the list of genes and their parts (list or range of exon and/or codon numbers); (2) multi-step primer design that allows efficient optimizing of primer design parameters (GC-content, Tm, amplicon length etc.); (3) checking primers for non-target hybridization in genome; (4) checking primers for overlapping with variable genome sites, i.e. SNPs; (5) designing primers for nested PCR; (6) designing primers for anchored PCR when target region is amplified from one sequence-specific primer and one primer that is complementary to an adapter sequence; (7) designing primers for whole defined genome region; (8) automatic distribution of primer pairs between multiplex reactions based on secondary structures and non-target amplicons that can be formed by primers from different pairs.

## 2 Materials and Methods

NGS-PrimerPlex code was written in Python using free-available Python-modules (**Supplementary Table 2**) and BWA program (Li and Durbin, 2009). It can be run as a standalone program (only for MacOS and Linux users) or inside docker image with or without GUI (for users of any OS, **Supplementary Figure 1**). Docker image allows someone not to install and not to download additional files but use it immediately after downloading the main package. NGS-PrimerPlex and its manual are available at GitHub server (https://github.com/aakechin/NGS-PrimerPlex).

### 2.1 Getting genome coordinates based on a list of genes and their parts

NGS-PrimerPlex contains script getGeneRegions.py that reads the table of genes from the input file, for which chromosome is determined from CSV-file that is created automatically from GenBank-files, and exon/codon coordinates are extracted from the corresponding GenBank-file. User can specify numbers of exons or codons which are necessary to be included into the targeted NGS-panel developed. Human reference genome versions hg19 or hg38 or reference genome of other organisms can be used. Output file can be used in the next script for primer design NGS-primerplex.py. Whole NGS-PrimerPlex package functionalities are reflected in **Figure 1.**

**Figure 1.**
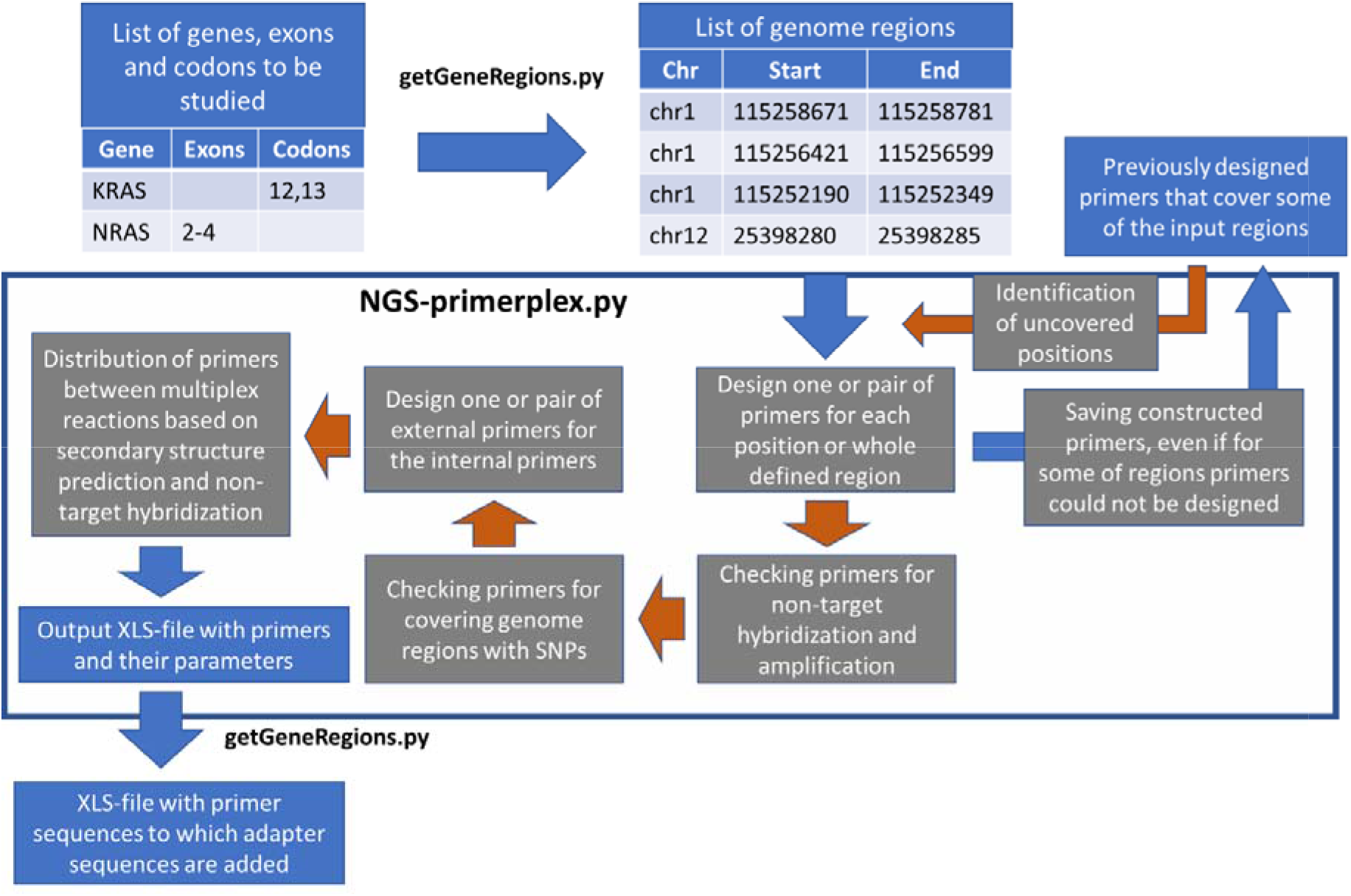
Whole workflow of NGS-PrimerPlex. Blue arrows denote obligatory steps; arrows filled with brown denotes optional steps that can be applied by user. Blue rectangles denote input and output files; grey rectangles denote steps of the whole workflow.

### 2.2 Multi-step primer design

By default, NGS-PrimerPlex splits input genome regions onto distinct positions and tries to construct primers for each of them using primer3-py that includes primer3 library (Untergasser et al., 2012). For each position, the program tries to design three types of primers: so that the right primer was close to the studied position, the left primer was close to the studied position, and without any of these restrictions (**Supplementary Figure 2**). This is necessary for subsequent successful joining of primers into a set of primer pairs that amplify a single extended genome region (e.g. whole exon). Unlike other programs (e.g. hi-plex) which design primer pairs not overlapping at all, NGS-PrimerPlex allows overlapping but considers it during distribution into multiplex reactions. This increases chances of successful primer design, particularly for genome regions with complex structure. At the same time, NGS-PrimerPlex allows user to forbid splitting of some regions, e.g. during primer design to detect EGFR exon 19 deletion (Oskina et al., 2017).

Another feature of NGS-PrimerPlex that may be useful for complex genome regions is a multi-step primer design. Frequently, existing tools suggest user to choose parameters of primer design and at the end of the design process program can give an error that primers could not be constructed with the defined parameters. After choosing less strict parameters, user has to start the process from scratch that significantly slows down parameter optimization. NGS-PrimerPlex saves primers that could be designed with more strict parameters and constructs primers only for other regions. On the other hand, this feature can be used to create personalized NGS-panels and expanding existing ones that are targeted for patient-specific genome regions, when primers for some of these regions have been obtained earlier that can not be done with other programs.

### 2.3 Non-target hybridization in genome and overlapping with SNPs

On-target is one of the main characteristics of targeted NGS-panels (Samorodnitsky et al., 2015) and it depends on whether the primers hybridize to non-target genome regions, including with substitutions/insertions/deletions as well as with other primers from the same multiplex reaction. For doing it, NGS-PrimerPlex uses BWA program that can readily find all regions to which a nucleotide sequence can be mapped. For the similar regions (but not necessarily identical ones, because the user can search for primer targets with mismatches), the program checks if primer has the same last two nucleotides from 3’-end as non-target region, that is not performed by existing tools (e.g. MPD and hi-plex). This allows identifying regions that will really give non-target amplicons but not only has homology with primers. Comparing genome coordinates for different primers, NGS-PrimerPlex filters out primer pairs that can lead to non-target amplification in multiplex reaction.

Another significant characteristic of NGS-panels is a uniformity of coverage (Samorodnitsky et al., 2015) which depends, among other things, on the sequence homogeneity of the sequence complementary to primer among different samples. Therefore, NGS-PrimerPlex checks if any of the primers designed overlaps with SNPs from the dbSNP database (Sherry et al., 2001) using pysam Python-module and dbSNP VCF-file. The user has an opportunity to check only part of primer for overlapping with SNP (e.g. only last 10 nucleotides that are especially important, as we showed in this work) and to define minimal frequency of SNP in population for which primers will be checked. Both features are unique for NGS-PrimerPlex.

### 2.4 Nested and anchored multiplex PCR

Nested multiplex PCR is applied in many areas of molecular biology where higher specificity is necessary, particularly during the detection of low-frequent somatic mutations in tumors (Zheng et al., 2014). It increases the specificity by selecting only amplicons which contain both internal and external primers (**Supplementary Figure 3**). NGS-PrimerPlex allows users to design primers for nested PCR with subsequent distribution of four primers among multiplex reactions considering both secondary structure and non-target product formation for internal and external primers.

Anchored multiplex PCR that applies one primer hybridizing to gene-specific region and one primer been complement to adapter sequence allows amplifying regions with unknown or partially highly variable sequence (**Supplementary Figure 3**), e.g. to detect gene fusion mutations without prior knowledge of the fusion partners (Schenk et al., 2017; Zheng et al., 2014). And NGS-PrimerPlex can also design primers for this type of multiplex PCR.

### 2.5 Distribution of primers among multiplex reactions

The final step of primer design is a distribution of the constructed primers among user defined number of multiplex reactions. NGS-PrimerPlex supports distribution of all primers into any number of multiplexes (unlike e.g. MPD that is capable of distributing into small multiplexes of about 2-15 primer pairs) as well as distribution of some regions into specific groups of multiplexes (e.g. when it is necessary to separate some genes into different multiplexes). The distribution is performed using networkx Python module which creates graph, edges of which mean two primer pairs not producing any secondary structures, not overlapping by the product, and not producing non-target amplicons. For joining primer pairs into set of primer pairs that amplify single extended genome region (e.g. whole exon), it searches for a shortest path from the start of such region to the end, both of which are also included into the graph. For the subsequent distribution, NGS-PrimerPlex tries to find clique in the constructed graph, i.e. such subset of vertices that every two distinct vertices in the clique are adjacent. Searching for clique is performed until all primer pairs are distributed.

### 2.6 NGS library preparation and sequencing

To confirm efficiency of NGS-panels designed with NGS-PrimerPlex tool, we applied it to sequence coding exons of *LRRK2* gene, associated with Parkinson’s disease (Di Maio et al., 2018). The *LRRK2* gene includes 51 exons, all of which are protein-coding (7784 bp including two intron nucleotides near exon-intron junctions). DNA was extracted from the peripheral blood leukocytes as it was described earlier (Kechin et al., 2018). NGS libraries were prepared with in-house amplicon-based approach using two-step amplification: (1) enrichment of target regions; (2) inclusion of adaptors. The libraries were sequenced with MiniSeq High Output kit (300 cycles). NGS-reads were analyzed with workflow that is similar to BRCA-analyzer’s one (Kechin et al., 2018). NGS-reads were trimmed with Trimmomatic (Bolger et al., 2014) and mapped to human reference genome (hg19). Coverage was evaluated with samtools (Li et al., 2009) and Python-scripts. Variations were called with Pisces (https://github.com/Illumina/Pisces).

## 3 Results

To evaluate throughput of NGS-PrimerPlex, we applied it to design primers for coding sequences of several sets of genes that are studied in clinics with existing commercial NGS-panels (**Supplementary Table 3**). For genes from Tumor15 panel, we used whole coding sequences of these genes regardless of which parts of them are clinically significant. We showed that NGS-PrimerPlex can design NGS-panels for a sizable number of genes and genome positions in reasonable time on computers with different performances.

### 3.1 Wet-lab validation

117 primer pairs were automatically designed without checking primers for covering SNPs (to evaluate their effect depending on remoteness from 3’-end of primer) and sorted to three multiplex reactions (each of 39 primer pairs) with NGS-PrimerPlex to sequence coding regions of the *LRRK2* gene (**Supplementary File 1**). Totally, 403 DNA samples from 403 patients were analyzed with the created NGS-panel. For samples with a median coverage of target regions by at least 30 reads (354 samples), median percent of targets covered by at least 30 reads was 97.4% (**Figure 2**).

**Figure 2.**
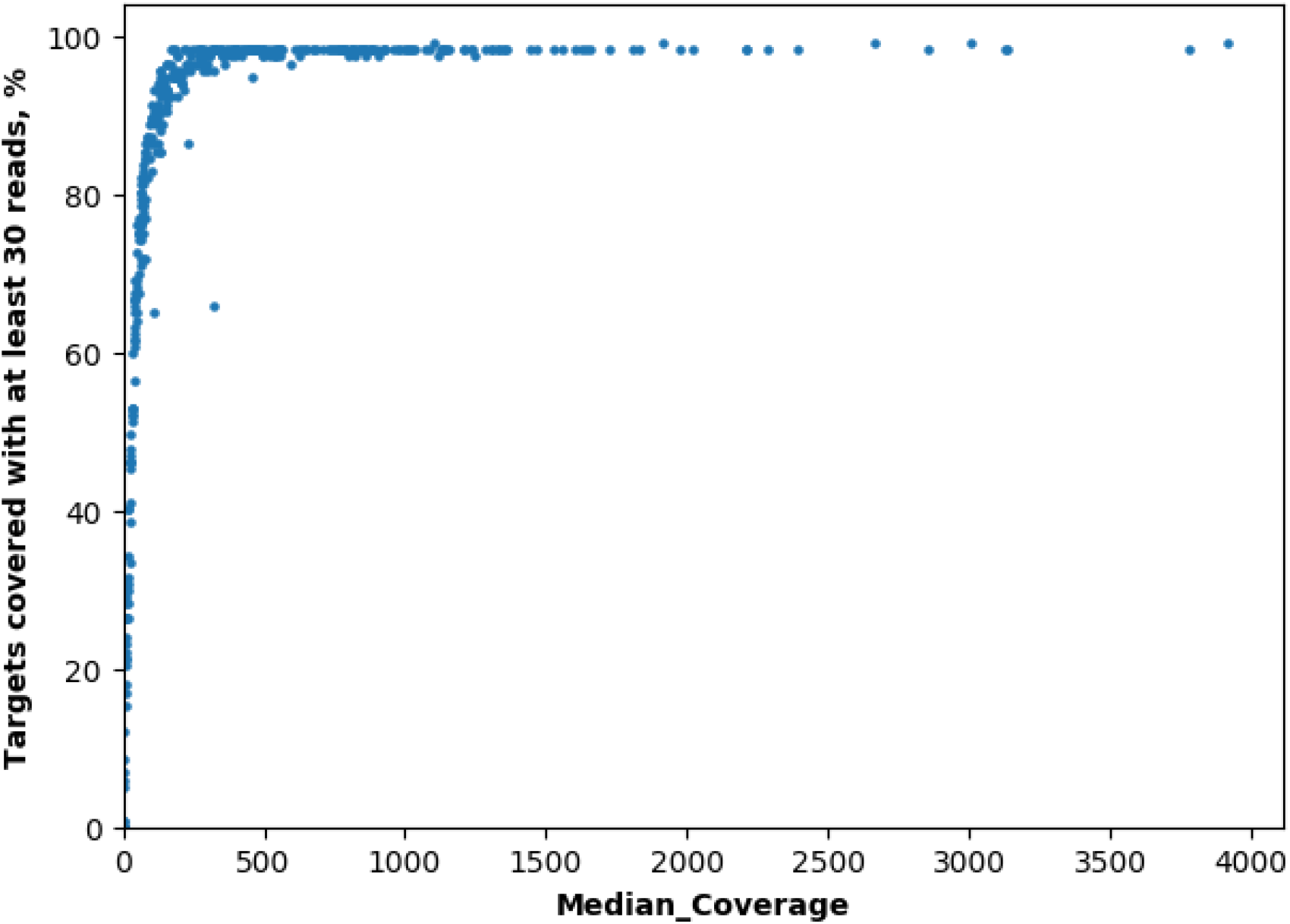
Percent of targets covered by at least 30 reads vs median coverage for samples sequenced for *LRRK2* gene. Each dot is a distinct DNA sample.

For other samples, median coverage of target regions was less than 30, likely due to low amounts or low quality of DNA used. One amplicon was absolutely uncovered, one amplicon was covered by at least 30 reads only for five samples, three amplicons had a high variation in coverage between samples due to overlapping with SNPs (rs73097447 located at 2^nd^ nucleotide of primer from 3’-end: 68% of reference allele vs 32% of alternative allele, rs1429478461 located at 4^th^ nucleotide of primer: 61% of reference allele vs 39% of alternative allele for hg19, and rs711176013 located at 8^th^ nucleotide of primer: 41% of reference allele vs 59% of alternative allele for hg19) (**Figure 3**). Thus, primers covering an SNP by −8 to −10 nucleotide from the 3’-end demonstrate a high variation in the amplification efficiency.

**Figure 3.**
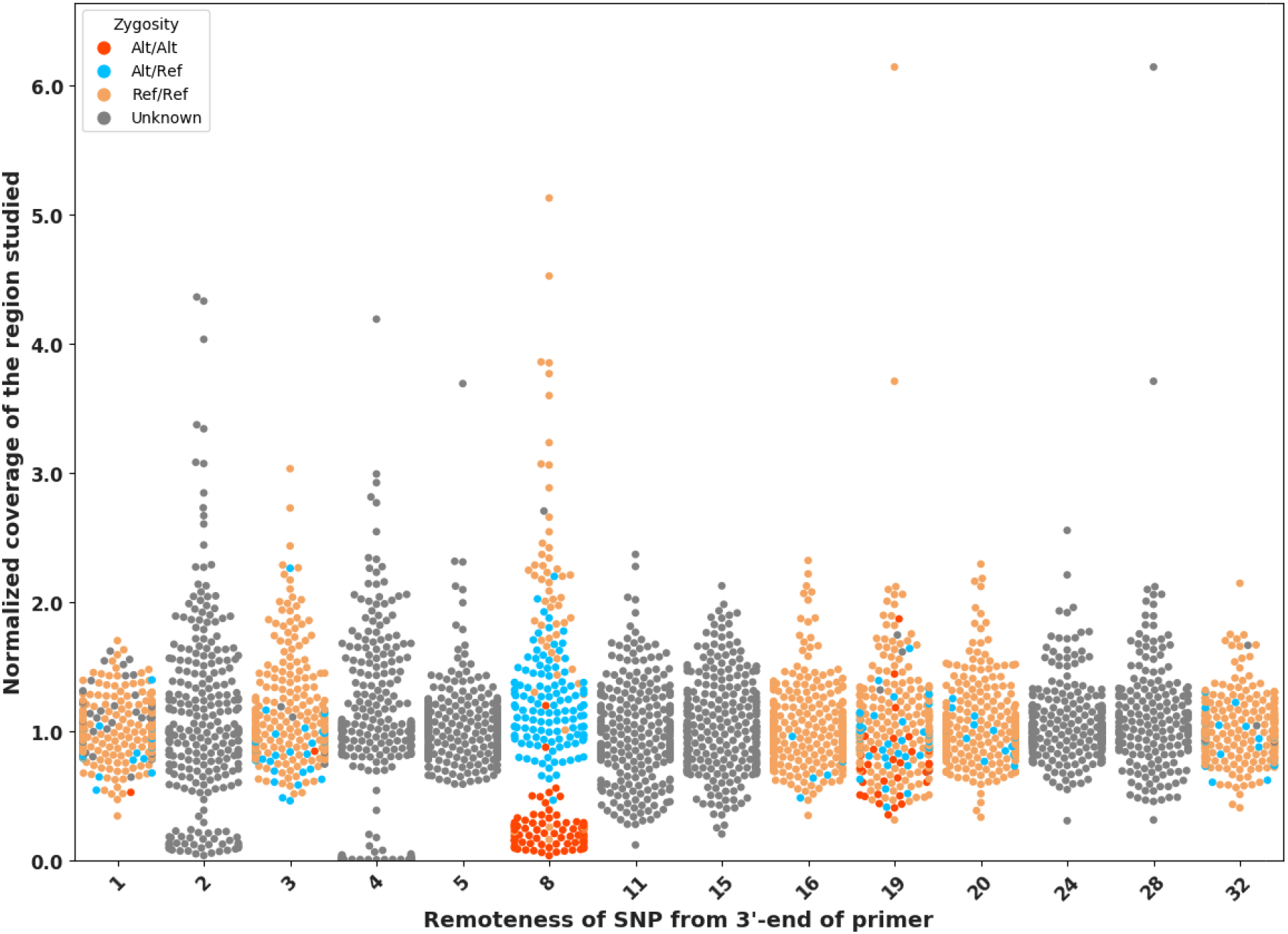
Effect of SNPs under primers on coverage values depending on its remoteness from 3’-end of primer. Each dot reflects double normalized coverage of a region studied depending on genotype of SNP under primer. Unknown means that SNP was located in intron region that was not covered by the NGS-panel, and its genotype could not be known. SNPs for the regions studied have the following frequencies in 1000Genomes database, respectively: (1) 1%, (2) 32%, (3) 6%, (4) 39%, (5) 59%, (8) 59%, (11) 59%, (15) 32%, (16) 0.1%, (19) 28%, (20) 10%, (24) 40%, (28) 59%, (32) 6%.

Commonly, we should avoid primers covering SNPs. And NGS-PrimerPlex certainly has an opportunity to turn it off by skipping -nucs argument. And then the program will not allow any SNPs under primers. However, this is not always possible, e.g. for regions that contain many SNPs and should be studied in the whole. In this case, we can limit checking for SNP overlapping to about 10 nucleotides from 3’-end and to SNPs with frequency more than 5%, that can be defined in NGS-PrimerPlex and is absent in other tools (e.g. MPD).

After this experiment, primers covering SNPs by at least one of 10 nucleotides from 3’-end of primers were removed and redesigned with NGS-PrimerPlex using -draft and -snps parameters and by increasing -primernum1 parameter, leaving other primers the same.

## 4 Discussion

NGS-PrimerPlex is a high-throughput cross-platform tool for automatic primer design for different multiplex PCR applications, including creation of new targeted NGS-panels. It includes many useful features that significantly facilitate primer design process. Among them, NGS-PrimerPlex can (1) automatically choose genome regions of targets, (2) expand and improve existing NGS-panels (3) has more accomplished checking for non-target hybridizations and covering SNP by primer, (4) design primers for nested and anchored multiplex PCR and (5) distribute primers among defined number of multiplex reactions regardless of primer pool sizes. All of the listed features are not supported or have limited use by existing tools for multiplex primer design.

## Acknowledgements

The study was supported under Russian State funded budget project of ICBFM SB RAS #AAAA-A17-117020210025-5 “Development of the methods of personalized medicine”.

## Conflict of Interest

none declared.

## Notes

### Competing Interest Statement

The authors have declared no competing interest.

