## Supplementary for "NGS-PrimerPlex: high-throughput primer design for multiplex polymerase chain reactions"

### Supplementary materials

#### Supplementary Table 1. Comparison of NGS-PrimerPlex and other tools functionality:

**a** – choosing genome regions to amplify by gene names and their parts.

**b** – primer design for all chosen genome regions.

**c** – checking primers for non-target hybridization and non-target amplification.

**d** – checking primers for covering variable genome sites that contain high-frequent SNPs.

**e** – automatic primer distribution for several multiplex reactions, considering secondary structures and non-target amplicons that can be formed by primers from different pairs.

**f** – primer design for nested PCR.

**g** – primer design for anchored PCR when target region is amplified from one sequence-specific primer and one primer complemented to adapter sequence. It can be useful for detection of gene fusions.

**h** – has graphical or web-interface

**yes/no** – means that this tool includes or does not include such functionality, respectively.

± – means that such functionality is partial (e.g. distribution among pools is restricted by maximum of 15 primer pairs in one pool)

**?** – means that such functionality was claimed but tool wasn’t acceptable

| **#** | **Tool** | **Target tasks** | **Reference** | **a** | **b** | **c** | **d** | **e** | **f** | **g** | **h** |
| --- | --- | --- | --- | --- | --- | --- | --- | --- | --- | --- | --- |
|  | **NGS-PrimerPlex** | Multiplex PCR, Targeted NGS, qPCR | - | **yes** | **yes** | **yes** | **yes** | **yes** | **yes** | **yes** | **yes** |
| 1 | Hi-Plex2 | Targeted NGS | (Hammet *et al.*, 2019) | no | **yes** | no | no | no | no | no | no |
| 2 | Multiplex Primer Design | Targeted NGS | (Wingo *et al.*, 2017) | no | **yes** | **yes** | **yes** | ± | no | no | ? |
| 3 | PrimerPooler | Targeted NGS | (Brown *et al.*, 2017) | no | no | no | no | **yes** | no | no | no |
| 4 | ThermoAlign | Targeted NGS | (Francis *et al.*, 2017) | no | **yes** | no | no | no | no | no | no |
| 5 | PrimerMapper | PCR | (O’Halloran, 2016) | no | **yes** | no | no | no | no | no | no |
| 6 | MRPrimerW | qPCR | (Kim *et al.*, 2016) | no | **yes** | no | no | no | no | no | yes |
| 7 | MSRE-HTPrimer | Epigenetics | (Pandey *et al.*, 2016) | no | **yes** | no | no | no | no | no | ? |
| 8 | HTP-OligoDesigner | Cloning | (Camilo *et al.*, 2016) | no | **yes** | no | no | no | no | no | yes |
| 9 | PrimerView | PCR, NGS? | (O’Halloran, 2015) | ? | ? | ? | ? | ? | ? | ? | no |
| 10 | PRIMEGENSw3 | PCR, epigenetics | (Kushwaha *et al.*, 2015) | no | **yes** | no | no | no | no | no | yes |
| 11 | Hiplex-Primer | Targeted NGS | (Nguyen-Dumont *et al.*, 2013) | no | **yes** | no | no | no | no | no | no |
| 12 | Optimus Primer | Targeted NGS | (Brown *et al.*, 2010) | **yes** | **yes** | no | no | no | no | no | yes |
| 13 | MPprimer | Multiplex PCR | (Shen *et al.*, 2010) | no | **yes** | no | no | no | no | no | ? |
| 14 | QuantPrime | qPCR | (Arvidsson *et al.*, 2008) | no | **yes** | no | no | no | no | no | ? |
| 15 | JCVI Primer Design Tool | Sanger Sequencing | (Li *et al.*, 2008) | **yes** | **yes** | no | no | no | no | no | no |
| 16 | BatchPrimer3 | PCR, Sanger Sequencing | (You *et al.*, 2008) | no | **yes** | no | no | no | no | no | yes |
| 17 | PrimerPlex ^*^ | Targeted NGS | http://www.premierbiosoft.com/primerplex/ | **yes** | **yes** | **yes** | no | no | no | no | yes |
| 18 | MultiPLX | Multiplex PCR | (Kaplinski and Remm, 2007) | no | no | no | no | **yes** | no | no | yes |

#### Supplementary Table 2. List of Python-modules used in NGS-PrimerPlex.

| **Python-module** | **Tasks solved with this module** | **Reference or link** |
| --- | --- | --- |
| biopython | working with nucleotide sequencing | (Cock *et al.*, 2009) |
| argparse | reading input arguments | <https://docs.python.org/3/library/argparse.html> |
| primer3-py | primer design for one region or position | <https://github.com/libnano/primer3-py> and (Untergasser *et al.*, 2012) |
| pysam | working with SAM- and VCF-files | <https://github.com/pysam-developers/pysam> |
| xlrd | reading EXCEL-files | <https://pypi.org/project/xlrd/> |
| xlsxwriter | writing to EXCEL-files | <https://pypi.org/project/XlsxWriter/> |
| networkx | joining primer pairs to multiplex reactions | (Hagberg *et al.*, 2008) |
| numpy | choosing best combination of primer pairs | (van der Walt *et al.*, 2011) |

**Supplementary Figure 1.** Graphical interface of the NGS-PrimerPlex program. Four different windows are shown (from left top to right bottom): (1) main menu; (2) extraction of gene(s)’ CDS coordinates; (3) primer design; (4) settings. The most of settings have default values that can be used.

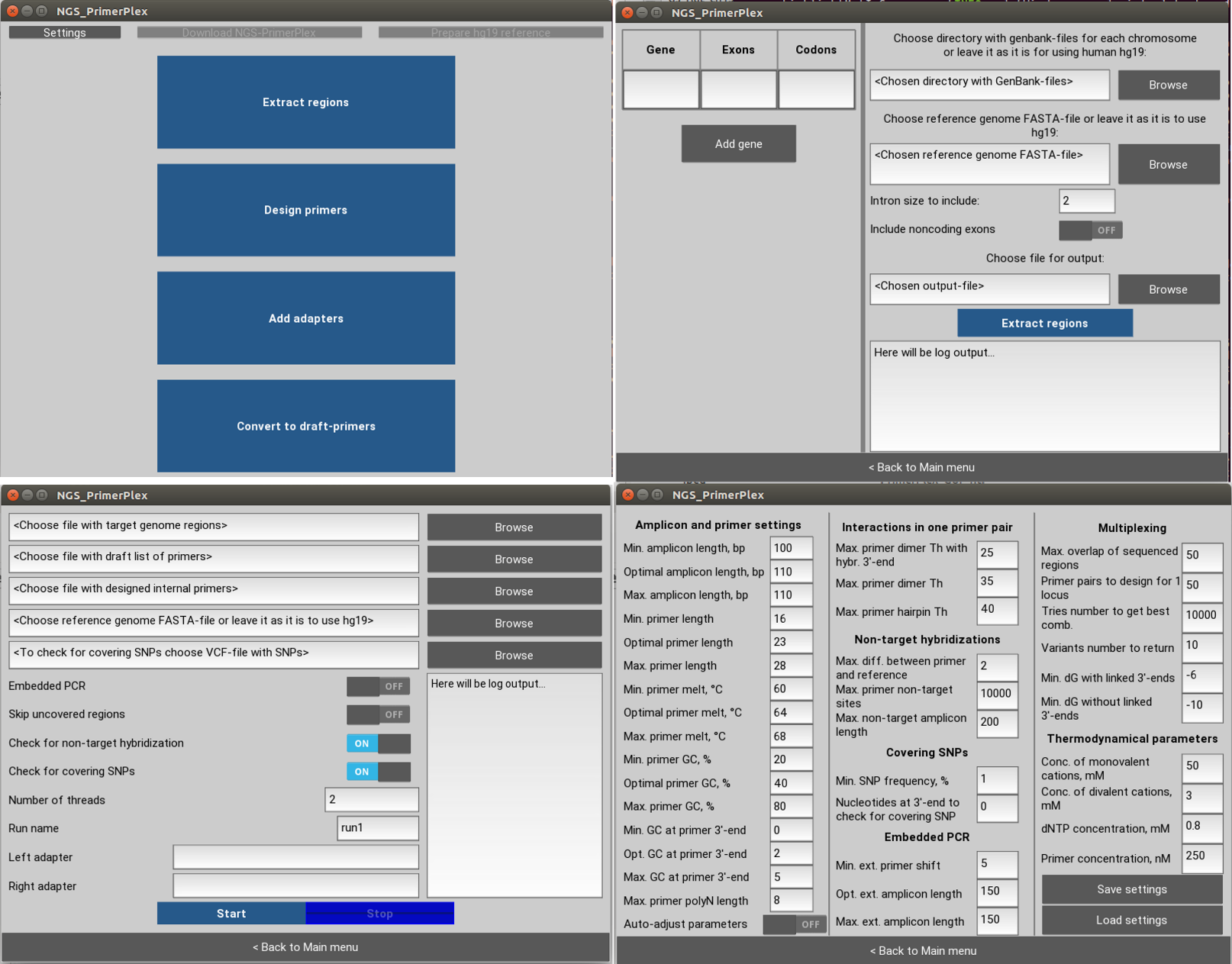

**Supplementary Figure 2.** (**A**) Approach used by NGS-PrimerPlex, when for each position, the program calls primer3 to design three types of primers: so that the right primer was close to the studied position, the left primer was close to the studied position, and without any of these restrictions. Such type of primer design gives more flexibility on the next steps, when primer pairs are combined into sets of primers that amplify whole studied region (e.g. exon). It is necessary because we can’t join two overlapping primer pairs into one multiplex reaction (**B**). x_i_, x_i+1_, x_i+2_ are genome positions placed one by one. L1.1 is a left primer of the first type for the first position; L1.2 is a left primer of the second type for the first position; R2.3 is a right primer of the third type for the second position etc. (**B**) Formation of non-target amplicon while joining of overlapping primer pairs into one multiplex reaction. At the same time, we need to design primers located one by one in order to read whole sequence of the studied region, because after sequencing, for fragment L2-R2, only part denoted with asterisk make sense for calling variants. More information about the primers in the amplicon-based targeted NGS you can read in (Kechin *et al.*, 2017).

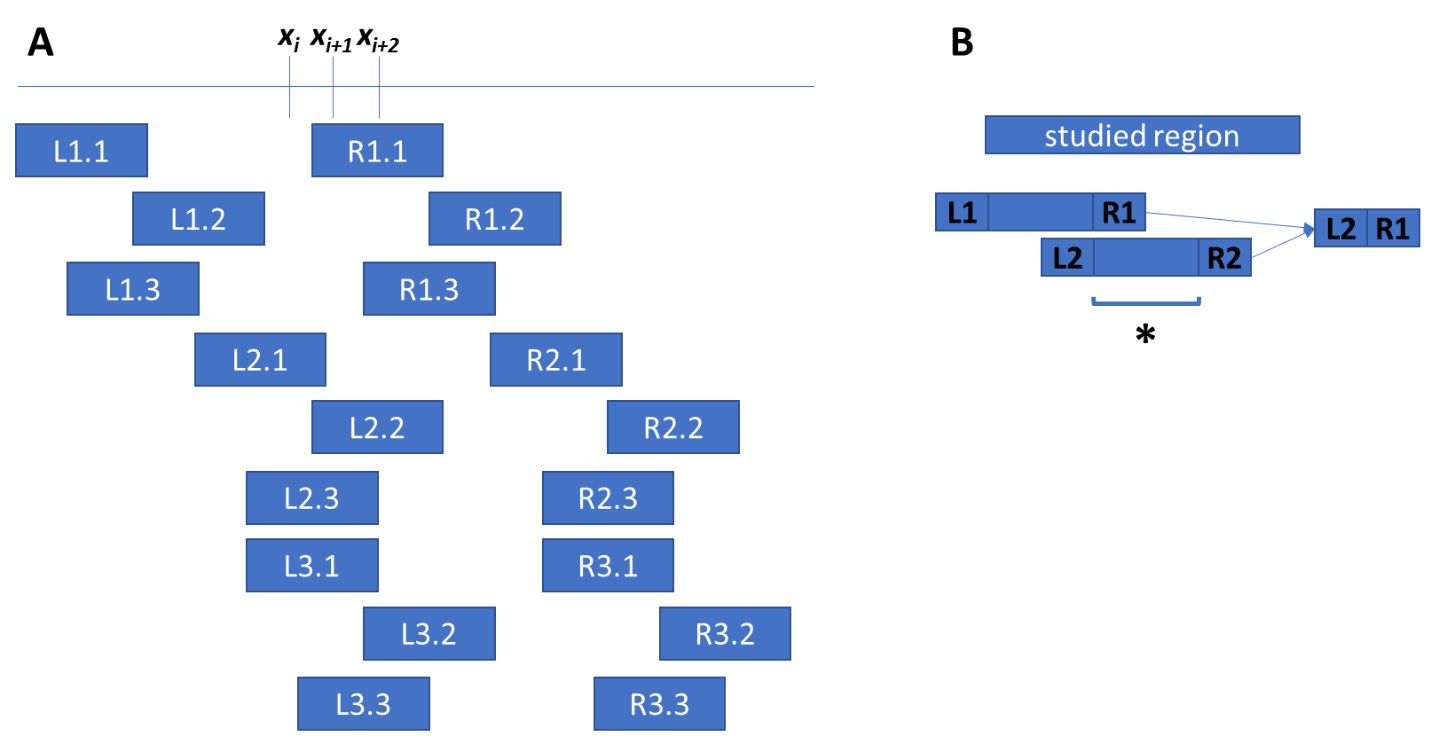

**Supplementary Figure 3.** The scheme of nested and anchored PCR. For nested PCR, brown arrows are external primers, dark blue arrows are internal primers. To design such four primers we should take into consideration the following conditions for external and internal primer: (1) non-target one primer hybridizations; (2) one primer pair non-target amplicons; (3) non-target amplicons for one pool primers (4) secondary structures between primers from the same and different amplicons. In case of primers with adapter sequences, we should take them into account while modeling secondary structures. For anchored PCR, we design one gene-specific primer that flanks highly variable region or region with unknown sequence (e.g. for gene fusions). We should consider (1) non-target one primer hybridizations and (2) secondary structures between one pool primers for different targets.

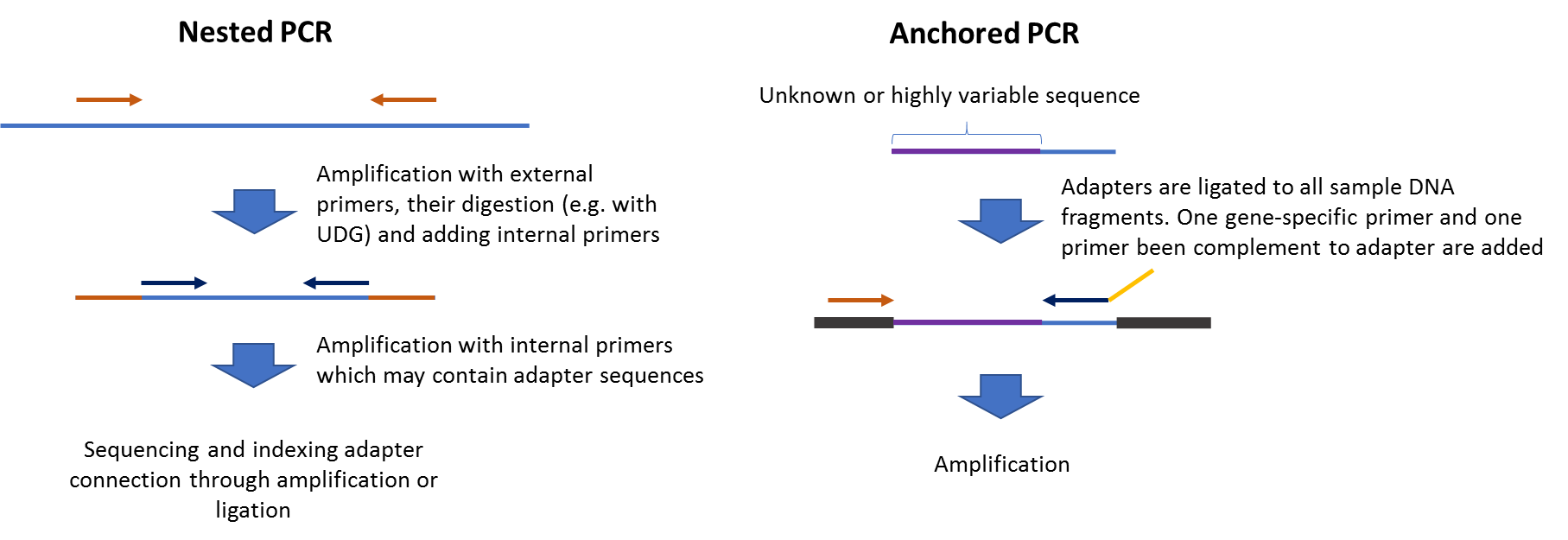

**Supplementary Table 3.** Time of primer design for clinically relevant regions that are already studied in some commercial NGS assays. All procedures were performed on three personal computers (1 – Intel Core i7-8700 3.2GHz, 64 GB RAM, Ubuntu 16.04; 2 – Intel Core i7-2700K 3.5GHz, 32 GB RAM Ubuntu 18.04; 3 – Intel Core i3-4130 3.4 GHz, 8 Гб, Windows 7 SP1 x64). Primer design was performed with checking for non-target hybridization and covering SNPs and distribution of primer pairs into multiplex reactions. The following parameters were defined (other were by default):

minimal – maximal (and optimal) amplicon length: 140–150 (150) or 80–130 (100) bp

minimal and maximal primer length: 16–35 nucleotides with optimal 25;

minimal and maximal melting temperature: 60–68 °C with optimal 64;

minimal and maximal GC-content of primers: 16–78 or 15–95 % with optimal 40 %;

minimal and maximal GC-content of the last 5 nucleotides of primers: 0–5 G/C-nucleotides;

maximal poly-N tract length: 8 or 9 bp;

mapping primers onto genome with allowing one error (substitution, insertion or deletion);

maximal overlap of neighboring amplicons: 10 or 50 bp

number of threads: 12, 8 or 2 depending on the computer used.

* – 11 bp of the GNA gene could not be covered due to the extremely high GC-content of this region (>95%).

| **List of genes and their parts** | **Examples of commercial NGS-assays (amplicons, multiplex reactions numbers)** | **Number of covered positions** | **Number of amplicons, their optimal length and multiplex reactions designed by NGS-PrimerPlex** | **Computer** | **Time of primer design, h** |
| --- | --- | --- | --- | --- | --- |
| Coding sequences of *BRCA1* and *BRCA2* genes | AmpliSeq™ BRCA Panel for Illumina® (265, 2)  Human *BRCA1* and *BRCA2* Panel Qiagen (?, 4) | 16037 | 208 (150 bp, 4 multiplexes) | 1 | 3.3 |
|  |  |  |  | 3 | 12.0 |
| *MLH1*, *MSH2*, *MSH6*, *PMS1*, *PMS2*, *POLE*, *MUTYH*, *PARP1*, *RECQL4* | Genes associated with hereditary forms of colorectal cancer | 30382 | 411 (150 bp, 6 multiplexes) | 1 | 12.4 |
| 15 genes that are commonly mutated in solid tumors (we used whole coding exons for all genes) | TruSight Tumor 15 (250, ?) | 34582-11=34571 * | 836 (100 bp, 10 multiplexes) | 2 | 12.1 |
